# *isoTarget*: a genetic method for analyzing the functional diversity of splicing isoforms *in vivo*

**DOI:** 10.1101/2020.04.30.070847

**Authors:** Hao Liu, Sarah Pizzano, Ruonan Li, Wenquan Zhao, Macy W. Veling, Yujia Hu, Limin Yang, Bing Ye

## Abstract

Protein isoforms generated by alternative splicing contribute to proteome diversity. Due to the lack of effective techniques, isoform-specific functions, expression, localization, and signaling mechanisms of endogenous proteins *in vivo* are unknown for most genes. Here we report a genetic method, termed *isoTarget*, for blocking the expression of a targeted isoform without affecting the other isoforms and for conditional tagging the targeted isoform for multi-level analyses in select cells. Applying *isoTarget* to two mutually exclusive isoforms of *Drosophila* Dscam, Dscam[TM1] and [TM2], we found that endogenous Dscam[TM1] is localized in dendrites while Dscam[TM2] is in both dendrites and axons. We demonstrate that the difference in subcellular localization between Dscam[TM1] and [TM2], rather than any difference in biochemical properties, leads to the two isoforms’ differential contributions to dendrite and axon development. Moreover, with *isoTarget*, we discovered that the subcellular enrichment of functional partners results in a DLK/Wallenda-Dscam[TM2]-Dock signaling cascade specifically in axons. *isoTarget* is an effective technique for studying how alternative splicing enhances proteome complexity.

## INTRODUCTION

Alternative splicing is a fundamental biological process that expands proteome diversity in eukaryotes. Genome-wide transcriptome analyses have shown that 90%-95% of human genes encode two or more isoforms (Baralle and Giudice, 2017). The percentage of multi-exonic genes that undergo alternative splicing is estimated to be 63% in mouse, 45% in *Drosophila*, and 25% in *C. elegans* (Lee and Rio, 2015). Alternative splicing is regulated by the coordination of RNA-binding proteins, RNA polymerase II, and epigenetic modifications of DNA (Baralle and Giudice, 2017). Perturbations of RNA splicing cause neurodevelopmental, cardiovascular, and other diseases (Baralle and Giudice, 2017; Scotti and Swanson, 2016). Despite these important findings, how distinct protein isoforms resulted from alternative splicing differ in their functions and regulations are poorly understood. In fact, the cellular functions, endogenous expression and localization, and signaling cascades of individual splicing isoforms are only known for a very small number of genes (Baralle and Giudice, 2017).

Protein isoforms encoded by alternative exons often differ in their structures and biochemical properties, which lead to distinct functions of the isoforms (Kelemen et al., 2013). In addition, different types of cells may express different splicing variants (Baralle and Giudice, 2017), which further diversifies the biological functions of different isoforms. Intriguingly, different protein isoforms of some genes are localized to distinct subcellular compartments within the same cell (Baralle and Giudice, 2017; Kelemen et al., 2013; Lee et al., 2016; Lerch et al., 2012; Yap and Makeyev, 2016). Compared to our understanding of how biochemical and expressional differences contribute to distinct functions of splicing isoforms, much less is known about whether and how isoform-specific subcellular localization contributes to distinct cellular functions. This is a challenging problem because solving it requires manipulating a specific isoform at its endogenous locus—without affecting other isoforms—in a cell-specific fashion.

Transgene-mediated overexpression of splicing variants of interest is widely used for studying isoform-specific functions and subcellular localization in specific cells. However, it is well documented that overexpressed proteins often do not mimic the endogenous proteins in their spatiotemporal expression, localization, and functions (Baralle and Giudice, 2017; Kelemen et al., 2013; Moriya, 2015; Prelich, 2012).

Here, we report a genetic method, termed *isoTarget*, for studying isoform-specific function, localization, and signaling of endogenous splicing isoforms of interest. This method allows to knock out a select isoform for functional studies and to tag conditionally the endogenous proteins for multi-disciplinary analyses in specific cells without affecting other isoforms. To achieve these, we created a <u>translational</u> stop sequence, used it to generate a cleavable cassette that contains an epitope tag for conditional tagging, and inserted the cassette into the exon encoding the isoform of interest. As a proof-of-concept, we applied *isoTarget* to study two mutually exclusive isoforms of *Drosophila* <u>D</u>own <u>s</u>yndrome <u>c</u>ell <u>a</u>dhesion <u>m</u>olecule (Dscam), Dscam[TM1] and [TM2] (Schmucker et al., 2000; Wang et al., 2004; Zhan et al., 2004). We report isoform-specific functions for Dscam[TM1] and [TM2] resulting from the distinct endogenous subcellular localization patterns of these two isoforms. We further describe a compartment-specific signaling pathway in the axon terminals, which involves Dscam[TM2], but not [TM1], as a result of the differential localization of the two isoforms and their functional partners. These findings illustrate the versatility of *isoTarget* in isoform studies and its effectiveness in uncovering mechanisms governing the expansion of proteome diversity by alternative splicing *in vivo*. In addition, they establish the causality between the subcellular localization and cellular function of Dscam splicing isoforms, demonstrating the critical role of subcellular localization in expanding the functional diversity of splicing isoforms.

## RESULTS

### The design of *isoTarget*

The transcriptional stop cassettes commonly used for conditional knockouts or knockins (Lakso et al., 1992) do not specifically disrupt the expression a select splicing isoform and are hence not applicable for isoform-specific studies. This is because RNA splicing occurs after transcription and, as a consequence, transcriptional stop cassettes disrupts the expression of all isoforms downstream of the targeted isoform (Figure 1A). To meet this challenge, we created a *translational stop* (*tlstop*) sequence by introducing multiple stop codons (TAA, TAG or TGA) into the DNA sequence encoding a non-catalytic region of β-Galactosidase (β-Gal) (Figures 1A, S1A). If the *tlstop* is present in an isoform-specific exon, it would lead to isoform-specific truncation during mRNA translation (Figure 1A).

**Figure 1.**
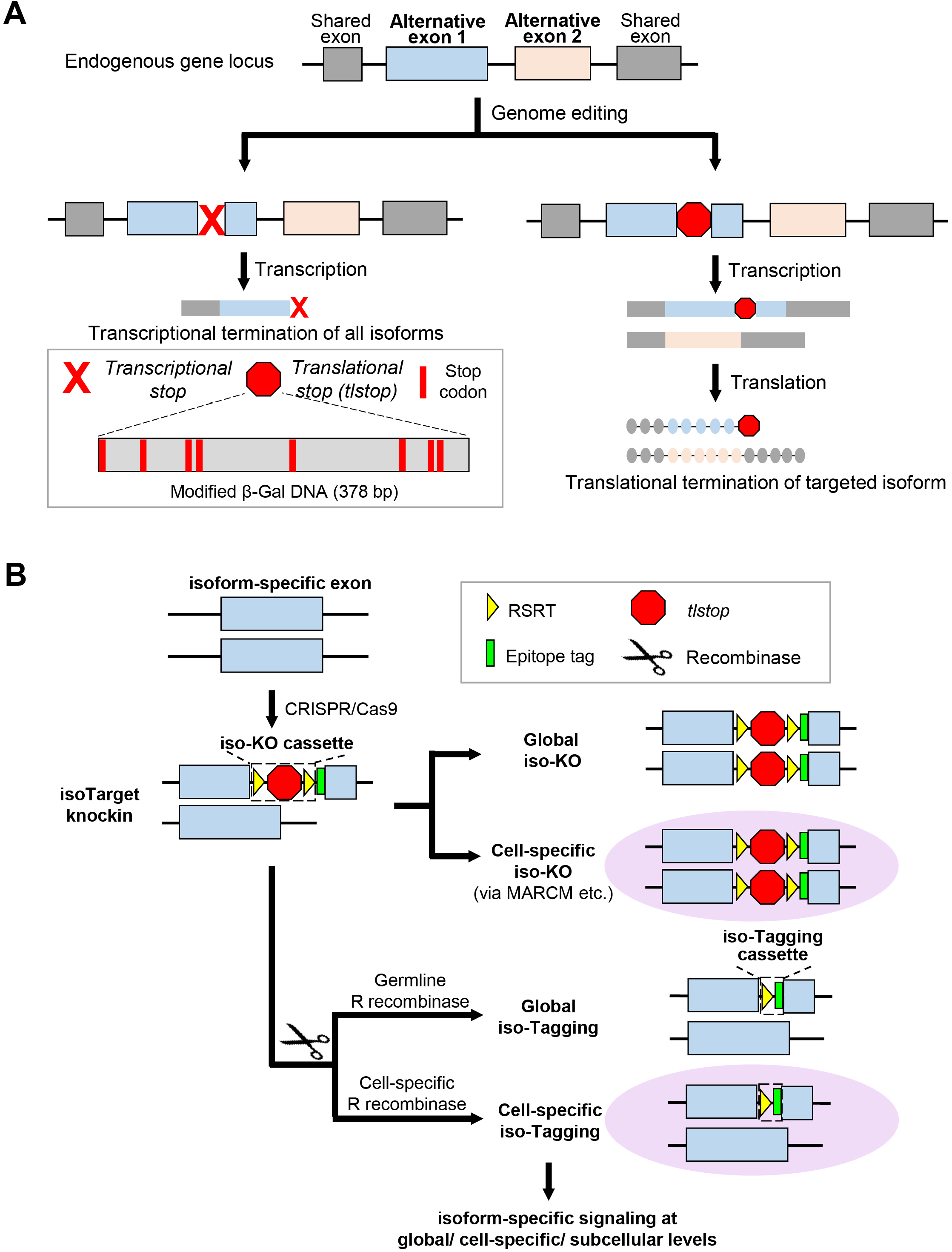
Design of *isoTarget* and its application to studying the functional diversity of splicing isoforms. (**A**) Introducing translational stops into alternative exons allows isoform-specific manipulations of the gene. Inserting the commonly used transcriptional stop cassette into alternative exon leads to the transcriptional termination of all isoforms downstream of the targeted exon (left branch). In order to truncate the targeted isoform, but not the transcription or translation of the other isoforms (right branch), we engineered a *translational stop* (*tlstop*) cassette by introducing multiple stop codons into the DNA sequence encoding a non-catalytic region of β-Gal (bottom left). (**B**) Combining *tlstop* with other genetic methods for multi-purpose studies of the targeted splicing isoform. The *isoTarget* cassette, which consists of an RSRT site, a translational stop (*tlstop*), another RSRT site, and an epitope tag, is inserted into the targeted isoform exon by CRISPR/Cas9-mediated genome editing. The insertion generates a loss-of-function allele of either the targeted isoform or the entire gene, depending on the length of the *tlstop* cassette (see Figure S3F). Single cells that are homozygous for the targeted allele can be produced by genetic mosaic techniques, such as MARCM. The endogenous isoforms can be visualized in specific neurons by selective expression of R recombinase to remove the RSRT-*tlstop*-RSRT cassette. Expression of R recombinase in female germline cells leads to tagging of the isoform in all cells that express this isoform in the progeny (“global iso-tagging”). Expression of R recombinase in specific cell types or single cells leads to tagging of the isoform in those cells. In the iso-Tagged flies, upstream regulators and downstream effectors of specific isoforms can be identified through genetic, cell biological, and biochemical analyses.

To achieve cell-type-specific labeling of targeted isoforms, the *tlstop* sequence is flanked by two R recombinase recognition sites (RSRT) followed by an epitope tag (Chen et al., 2014; Nern et al., 2011). When R recombinase is expressed to remove *tlstop*, the epitope tag is inserted in-frame into the targeted isoform, allowing the detection and biochemical analyses of endogenous proteins in cells of interest (Figure 1B). We refer to the RSRT-*tlstop*-RSRT as “iso-KO cassette” (for “*isoTarget* knockout cassette”), as its insertion into an isoform-specific exon is expected to create a loss-of-function mutant of this particular isoform. In *Drosophila*, the *iso-KO* alleles can be used in combination with the mosaic analysis with a repressible cell marker (MARCM) (Lee and Luo, 1999) to study isoform-specific function in targeted single neurons (Figure 1B). We refer to the RSRT-epitope resulted from the excision of the *tlstop* sequence as “iso-Tagging cassette”. As we show below, the iso-Tagging approach allows the investigation of upstream and downstream signaling mechanisms that involve targeted isoform at the organismal, cell-type-specific, or subcellular levels (Figure 1B).

### The validation of *isoTarget* and mitigation of off-target effects of translational-stop cassettes

To fulfill the designed applications, the *isoTarget* technique should meet the following requirements. First, the iso-KO cassette must abolish the function of the targeted isoform. Second, the iso-KO or iso-Tagging cassette in one isoform should not impair the expression of other isoforms. Third, iso-Tagging—resulted from the excision of the iso-KO cassette—should restore the isoform functions that are disrupted by the iso-KO cassette. We determined whether *isoTarget* met these prerequisites by testing on the *Drosophila Dscam* gene.

In *Drosophila*, exon 17.1 and 17.2 of *Dscam* gene encode two different transmembrane/juxtamembrane domains (Schmucker et al., 2000). Alternative splicing of these two mutually exclusive exons produces two isoforms called Dscam[TM1] and Dscam[TM2] (Figure S1B). We inserted the iso-KO cassette into the juxtamembrane domain in exon 17.1 (TM1) and 17.2 (TM2) (Figures S1B-C). In homozygous *Dscam[TM2]* iso-KO (*Dscam[TM2]^iso-KO^*) larvae, the axon terminal growth was dramatically impaired in the class IV dendritic arborization (C4da) neurons (Figures 2A-B, and H), a widely used model for studying dendrite and axon development (Grueber and Jan, 2004; Grueber et al., 2007; Jan and Jan, 2010; Ye et al., 2007). This is consistent with previous reports that *Dscam* is required for axon terminal growth in C4da neurons (Kim et al., 2013), and suggests that iso-KO cassette abolishes *Dscam[TM2]* functions. The impaired growth of axon terminals was completely rescued in homozygous global Dscam[TM2] iso-Tagging (*Dscam[TM2]^iso-Tagging^*) larvae resulted from the excision of the iso-KO cassette (Figures S1C, S2A-B), suggesting that tagging endogenous Dscam[TM2] with the epitope tag (V5) does not disrupt the function of the isoform.

**Figure 2.**
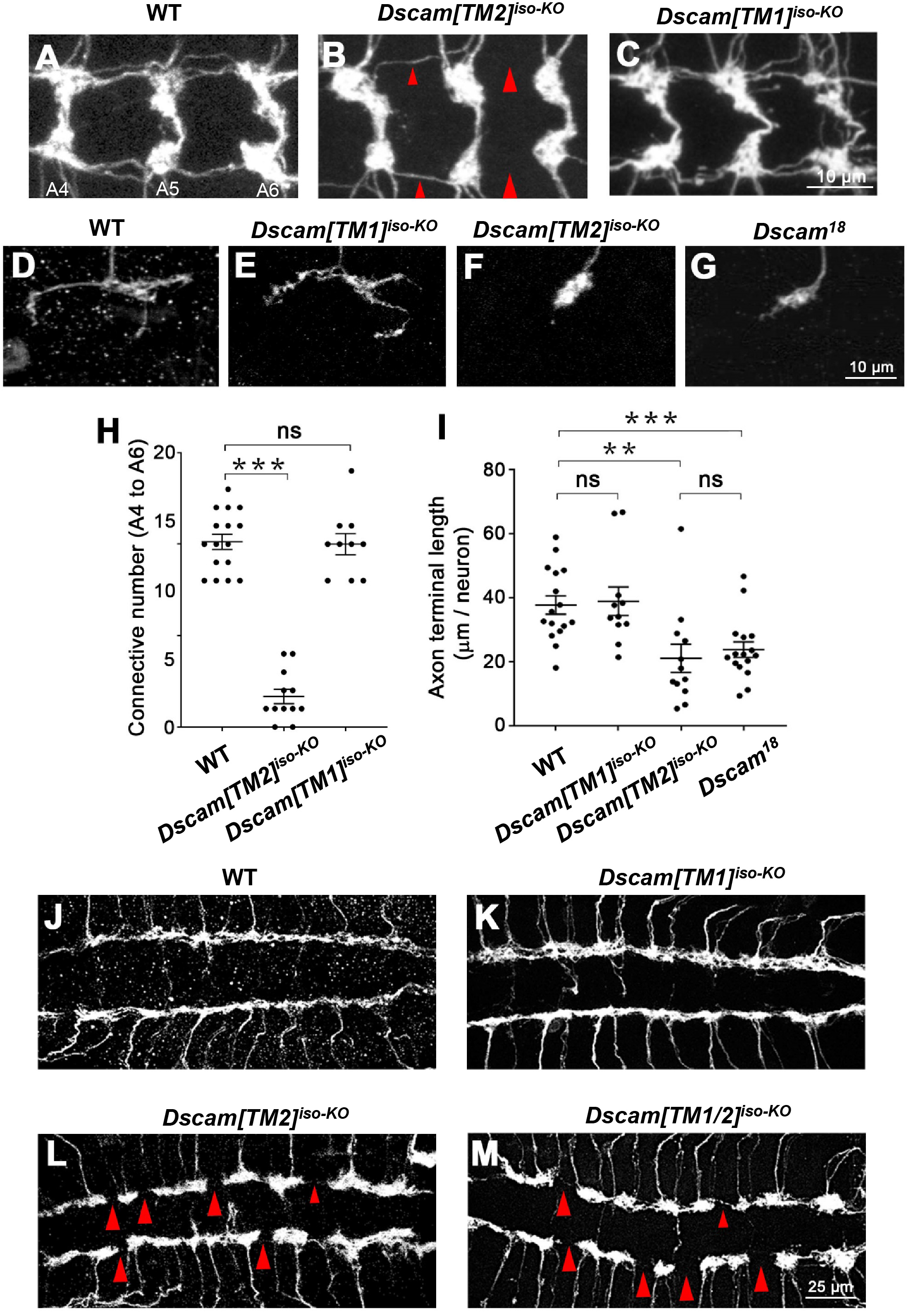
*IsoTarget* uncovers a specific role for the Dscam[TM2] isoform in axon terminal growth. (**A-C**) Compared with WT (**A**), global *Dscam[TM2]^iso-KO^* (**B**), but not *Dscam[TM1]^iso-KO^* (**C**), impairs axon terminal growth in C4da neurons. The C4da-specific driver *ppk*-Gal4 was used to label all C4da axon terminals in the CNS. Shown are representative images of abdominal segment 4 to 6 (A4-A6). The large red arrowheads point to the sites where longitudinal axon tracts are broken, and the small arrowheads point to where the tracts are thinned. (**D-G**) *Dscam[TM2]*, but not *[TM1]*, is required for the growth of axon terminals in single C4da neurons. The MARCM technique was used to generate single GFP-labeled C4da neurons that were homozygous of the indicated alleles. *Dscam[TM1]^iso-KO^* has no effect on axon terminal growth (**D** & **E**), while *Dscam[TM2]^iso-KO^* reduced the length of axon terminals to the same level as the loss of both isoforms (**F** & **G**). (**H**) Quantification of the number of axon connectives (i.e., the longitudinal branches) from A4 to A6. Unless specified otherwise, mean ± SEM is shown in all figures, and the statistical tests are one-way ANOVA followed by Student’s t test. *: p < 0.05; **: p < 0.01; ***: p < 0.001; ns: not significant (p > 0.05). (**I**) Quantification of presynaptic terminal length in the C4da neuron ddaC. (**J-M**) *Dscam[TM2]*, but not *[TM1]*, is required for the growth of axon terminals in C3da neurons. The longitudinal axon tracts of C3da neurons in the CNS remain intact in *Dscam[TM1]^iso-KO^* (**J** & **K**), but are disrupted in *Dscam[TM2]^iso-KO^* (**L**) or in mutants lacking both isoforms (**M**). The large red arrow heads point to the sites where longitudinal axon tracts are broken. The small red arrowheads point to the sites where longitudinal axon tracts are thinned.

Next, we determined whether *isoTarget* of one isoform affected the expression of another isoform by using quantitative real-time PCR and immunohistochemistry on *isoTarget* samples. *Dscam[TM1]* mRNA levels were not affected in the brains of homozygous *Dscam[TM2]^iso-KO^* or *Dscam[TM2]^iso-Tagging^* 3^rd^-instar larvae (Figure S2C). Unexpectedly, the [TM1] iso-KO cassette abolished *Dscam[TM2]* mRNA and protein expression, creating a *Dscam[TM1/2]^iso-KO^* mutant (Figures S3A and S3B-C). By contrast, the expression of *Dscam[TM2]* mRNA was not affected by *Dscam[TM1]* iso-Tagging cassette (Figure S3A). These results suggest that inserting a long piece of DNA in TM1-encoding exon disrupts the splicing of TM2-encoding exon of *Dscam* pre-mRNA (Figure S3F), possibly by overly extending the distance between exon 16 and the TM2-encoding exon (Anastassiou et al., 2006). To test this possibility, we reduced the size of the iso-KO cassette from 561 bp to 285 bp by cutting a 276 bp-fragment from the *tlstop* sequence. Strikingly, insertion of the short iso-KO cassette in [TM1] did not impair Dscam[TM2] expression, including both mRNA and protein expression (Figures S3A and S3D-E). Through these studies, we discovered that the impact of an *isoTarget* cassette on off-target isoforms can be mitigated by reducing the cassette length and found a new translational stop sequence for achieving this.

Taken together, these results validate the use of *isoTarget* for studying endogenous Dscam[TM1] and [TM2] isoforms.

### Uncovering the functions of Dscam isoforms in axon terminals with *isoTarget*

We previously demonstrated that in larval PNS neurons Dscam instructs the presynaptic terminal growth, which is a function that is distinct from Dscam’s role in neurite self-avoidance (Kim et al., 2013). However, whether both Dscam[TM1] and [TM2] contribute to this process is unknown. With *iso-KO* larvae, we found that knocking out the [TM2] isoform impaired axon terminal growth in larval C4da neurons (Figures 2A-B and H). By contrast, the axonal development remained intact in *Dscam[TM1]^iso-KO^* larvae (Figures 2C and H). Next, we combined *isoTarget* with the MARCM technique to determine whether the Dscam isoforms functioned cell-autonomously to regulate C4da axon terminal growth. Single C4da neurons that were homozygous for *Dscam[TM1]^iso-KO^* had normal axon terminal growth (Figures 2D-E and I). By contrast, loss of *Dscam[TM2]* in C4da neurons significantly impaired axon terminal growth to the same levels as loss of *Dscam* (Figures 2F-G and I).

Similar to C4da neurons, while C3da neurons in homozygous *Dscam[TM1]^iso-KO^* larvae showed normal growth of axon terminals, those in homozygous *Dscam[TM2]^iso-KO^* displayed dramatically reduced axon terminals growth, as evident in the gaps in longitudinal axon tracts (Figures 2J-L). The disruption in longitudinal axon tracts was also observed in *Dscam[TM1/2]^KO^* larvae (Figure 2M).

These results suggest that *Dscam[TM2]*, but not *Dscam[TM1]*, regulates the growth of axon terminals.

### Uncovering the functions of Dscam isoforms in dendrites with *isoTarget*

Dscam has been shown to mediate dendritic self-avoidance without affecting dendritic growth in the PNS neurons of *Drosophila* larvae (Hughes et al., 2007; Matthews et al., 2007; Soba et al., 2007). Again, it is unknown whether both Dscam[TM1] and [TM2] isoforms are responsible for this process. We applied *isoTarget* to answer this question. As expected from previous studies, homozygous *Dscam[TM1/2]^iso-KO^* mutant larvae, which lack both [TM1] and [TM2] functions, exhibited increased dendritic crossing in both C3da mechanosensors (Figures 3A-B and E) and C4da nociceptors (Figures S4A-B and E), indicating defective avoidance among dendrites of the same neuron. Different from what we observed in axons, neither *Dscam[TM1]^iso-KO^* nor *Dscam[TM2]^iso-KO^* caused any defect in dendritic self-avoidance in these neurons (Figures 3C-D and E, S4C-D and E).

**Figure 3.**
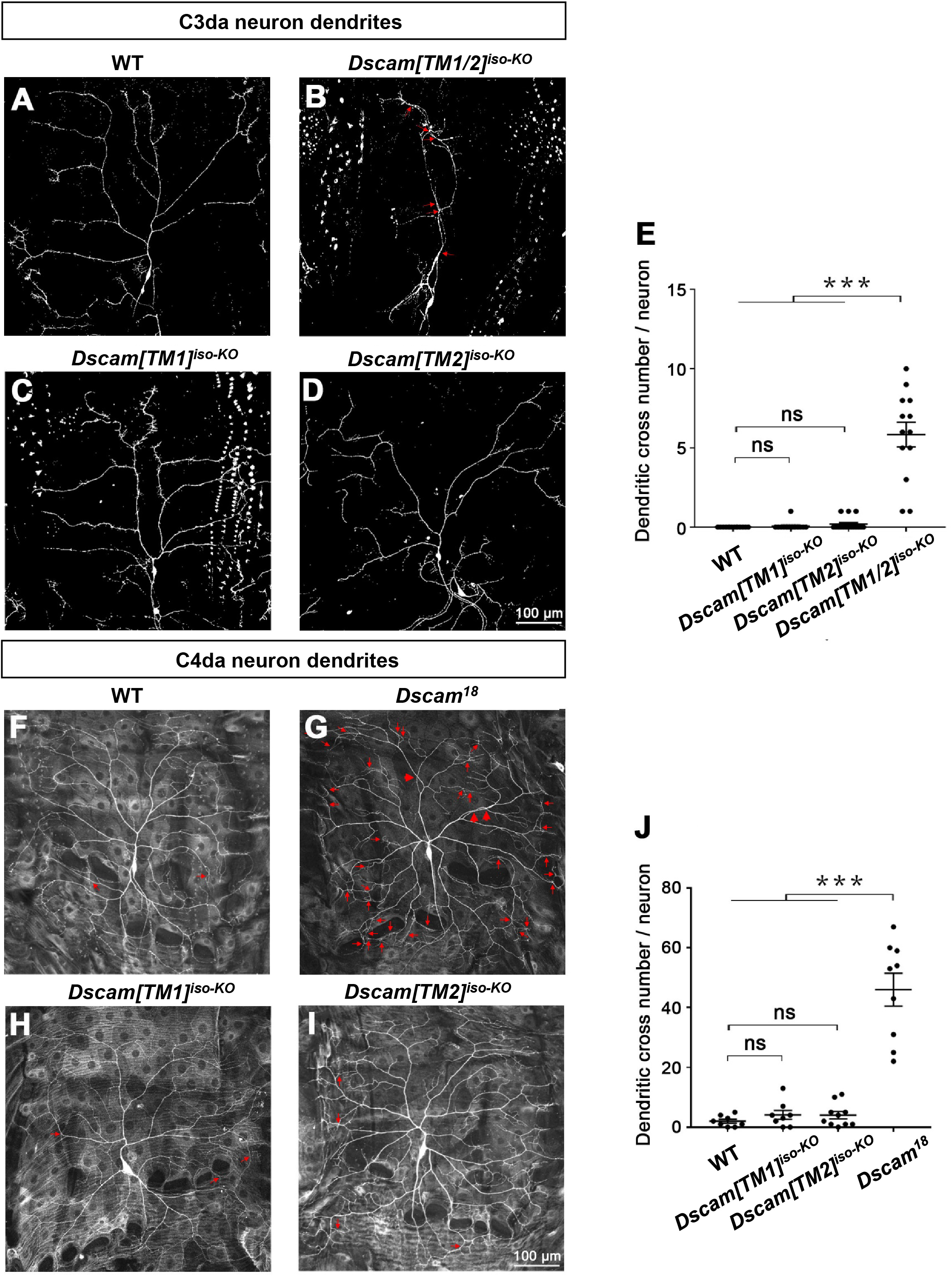
*IsoTarget* uncovers redundant functions for Dscam[TM1] and [TM2] in dendrite self-avoidance. (**A-D**) Dscam[TM1] and [TM2] function redundantly in mediating dendritic self-avoidance in C3da neuron. In 3^rd^ instar larvae, C3da dendrites rarely fasciculate or intersect with each other in wild-type (WT) (**A**), while loss of both *Dscam[TM1]* and *[TM2]* isoforms (*Dscam[TM1/2]^iso-KO^*) significantly impairs self-avoidance (**B**). Loss of either *Dscam[TM1]* (**C**) or *[TM2]* (**D**) does not affect dendritic self-avoidance. The red arrows point to dendritic crossing sites. (**E**) Quantification of dendritic branch crossings in the C3da neuron ddaF. (**F-I**) Dscam[TM1] and [TM2] function redundantly in mediating dendritic self-avoidance in C4da neurons. The MARCM technique was used. Small red arrows point to crossings of fine dendritic branches, and large red arrows point to crossings of major dendritic branches, which is only observed when both isoforms are lost. (**J**) Quantification of dendritic branch crosses in the C4da neuron ddaC.

We further combined MARCM with *isoTarget* to study cell-autonomous functions of targeted isoforms in single cells. As previously shown (Hughes et al., 2007; Matthews et al., 2007; Soba et al., 2007), single C4da neurons that were homozygous of *Dscam^18^*, which abolishes both [TM1] and [TM2] isoforms, showed significant self-avoidance defect (Figures 3G and J). By contrast, loss of either *Dscam[TM1]* or *[TM2]* in single C4da neurons did not cause any defect in dendritic self-avoidance (Figures 3H-I and J).

These results suggest that *Dscam[TM1]* and *[TM2]* function redundantly in dendritic self-avoidance. Thus, these two isoforms function differently in dendrite and axon development.

### Using *isoTarget* to identify the subcellular localizations of endogenous protein isoforms

Previous studies have shown that in CNS neurons transgenic Dscam[TM1] is restrained in somatodendritic compartments while [TM2] is in both dendrites and axons (Wang et al., 2004; Yang et al., 2008; Zhan et al., 2004). However, transgenic Dscam[TM1] and [TM2] were both found to be ubiquitously present in PNS neurons (Soba et al., 2007) (Figures 6B and D and data not shown). Does this discrepancy indicate that the subcellular localization of Dscam isoforms varies in different neuronal types? To answer this question, we applied iso-Tagging to examine the subcellular localizations of *endogenous* Dscam isoforms in both PNS and CNS neurons. We found that in PNS neurons both endogenous Dscam[TM1] and [TM2] were localized in the dendrites (Figures 4A-C). Interestingly, different from what we observed with *Dscam* transgenes, only endogenous Dscam[TM2] was localized in the presynaptic terminals of PNS neurons (Figures 4D and E). Similar isoform-specific localization patterns were observed in CNS neurons. By iso-Tagging in mushroom body (MB) neurons in the 3^rd^-instar larva, we observed both endogenous Dscam[TM1] and [TM2] in MB calyx (Figures 4F-H), which is a cluster of dendritic branches, but only [TM2] in the core fibers of axonal peduncles (Figures 4I-K). The finding that only [TM2] is localized in axons was further supported by the observation of Dscam proteins in axonal shafts that connect the PNS and CNS. In iso-Tagging larvae, despite a substantial amount of Dscam[TM1] signals in the neuropil region of the ventral nerve cord (VNC), no [TM1] signal was observed in axonal shafts (Figures S5A-B). By contrast, Dscam[TM2] puncta were abundant in axonal shafts in global *Dscam[TM2]^iso-Tagging^* larvae (Figure S5C). Taken together, we found that in PNS and CNS, both endogenous Dscam[TM1] and [TM2] isoforms are present in dendrites, while only [TM2] is in axons.

**Figure 4.**
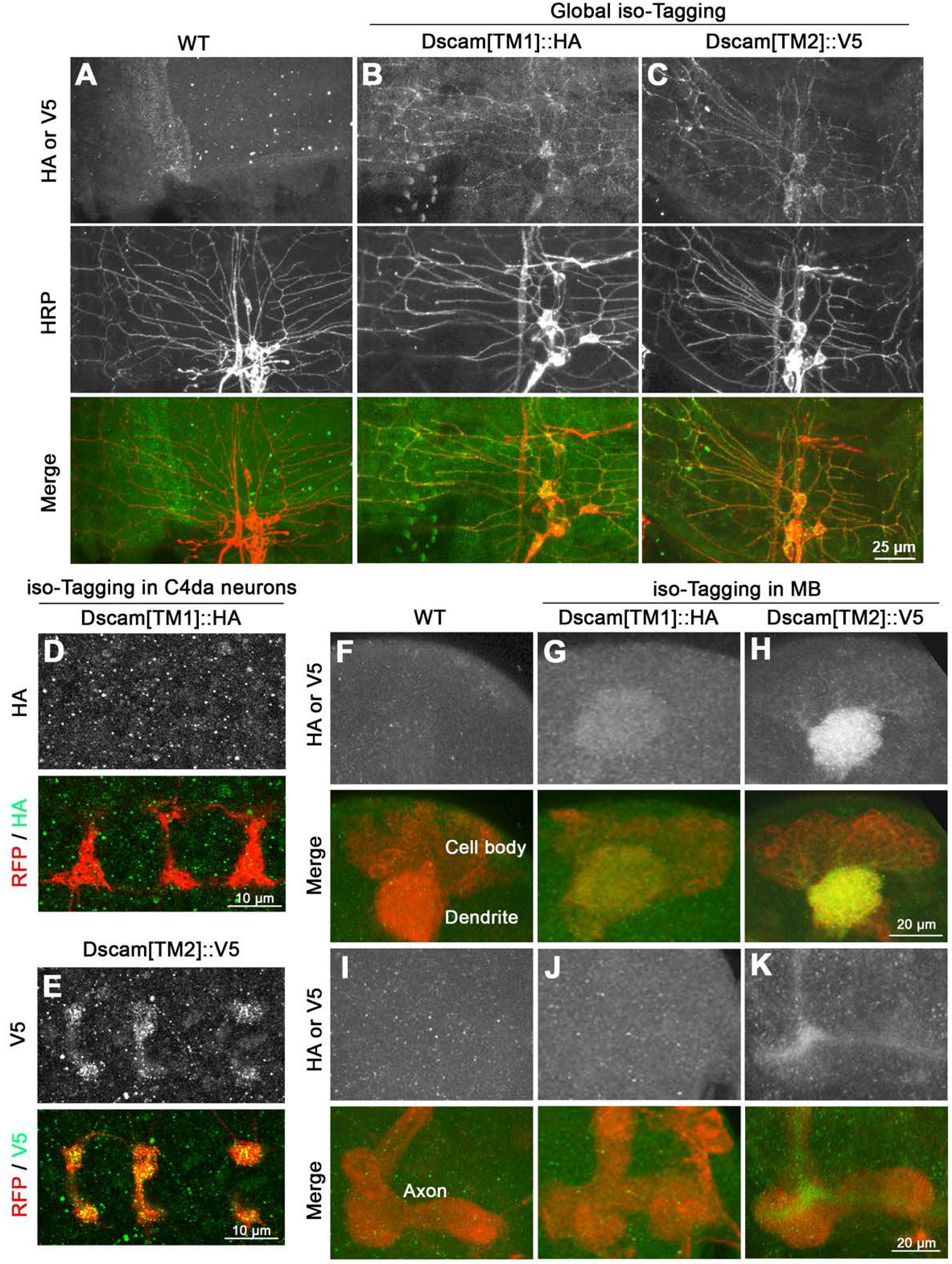
Using *isoTarget* to identify the subcellular localizations of endogenous Dscam isoforms. (**A-C**) Both Dscam[TM1] and [TM2] are localized in the dendrites of larval PNS neurons. At early 2^nd^ instar stage, compared with control (**A**), global iso-Tagging reveals the localization of endogenous [TM1] and [TM2] in the dendrites of PNS da neurons, which is double-labeled by the PNS neuron marker anti-HRP. (**D** & **E**) iso-Tagging shows that endogenous Dscam[TM2], but not [TM1], is in the axon terminals of C4da neurons in early 2^nd^ instar larvae. The C4da-specific driver *ppk*-Gal4 was used to tag endogenous Dscam[TM1] or [TM2] by driving the expression of UAS-R-recombinase and to label C4da axon terminals by driving the expression of UAS-mCD8::RFP. (**F-H**) Both Dscam[TM1] and [TM2] are localized in the dendrites of CNS neurons. Iso-Tagging in mushroom body neurons shows that both endogenous [TM1] and [TM2] are present in MB calyx, which is a cluster of dendrites in 3^rd^ instar larvae. The MB driver OK107-Gal4 was used to drive the expression of R recombinase for tagging endogenous Dscam[TM1] or [TM2] and the expression of mCD8::RFP for labeling MB morphology. (**I-K**) Dscam[TM2], but not [TM1], is localized in MB axons. Compared to the WT control, endogenous Dscam[TM2], but not [TM1], is detected in the axon peduncles of MB in 3^rd^ instar larvae.

**Figure 5.**
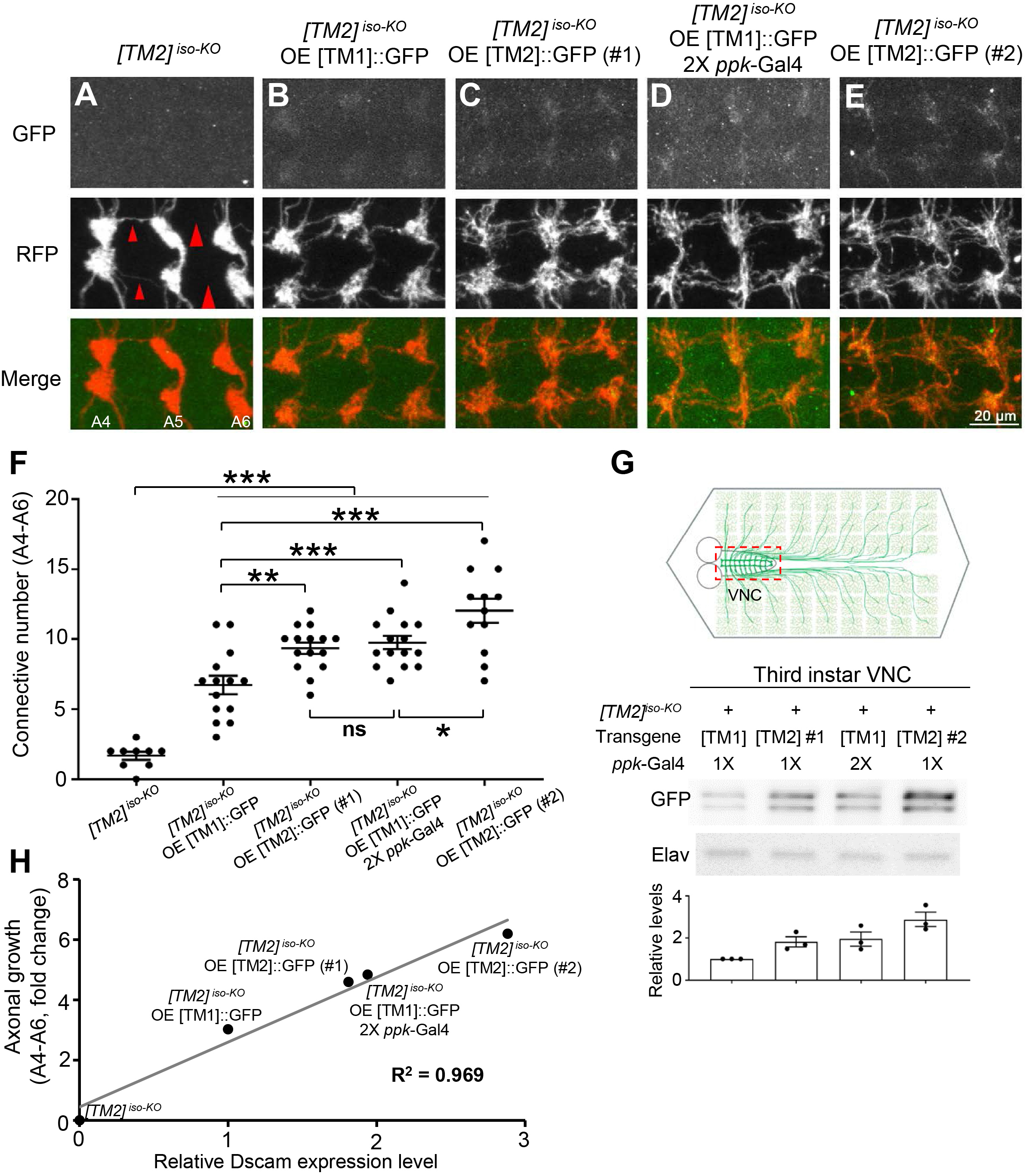
Dendrite-specific localization restrains endogenous Dscam[TM1] from functioning in axons. (**A**) *Dscam[TM2]^iso-KO^* dramatically impairs the axon terminal growth in C4da neurons. The large red arrowheads point to the sites where longitudinal axon tracts are broken, and the small arrowheads point to where the tracts are thinned. (**B**) Overexpression of a Dscam[TM1] transgene with a single copy of *ppk*-Gal4 leads to a low level of [TM1] in axon terminals and mitigates the axonal defect in [TM2] iso-KO. (**C**) Overexpression of the Dscam[TM2]#1 transgene with a single copy of *ppk*-Gal4 also mitigates the axonal defect in [TM2] iso-KO. (**D**) When the Dscam[TM1] transgene is driven by 2 copies of *ppk*-Gal4, it leads to higher levels of [TM1] in axon terminals than the overexpression driven by one copy of *ppk*-Gal4 and rescue effects that are comparable with [TM2] overexpression. (**E**) Overexpression of the Dscam[TM2]#2 transgene with a single copy of *ppk*-Gal4 leads to higher levels and rescue effects than that of [TM1] in C4da axon terminals. (**F**) Quantification of the number of C4da axon connectives in segments A4-A6. (**G**) Quantitation of transgenic Dscam::GFP levels in C4da axon terminals. The experiments were done in *Dscam[TM2]^iso-KO^* larvae. As shown in the schematic (top, green drawings), UAS-Dscam::GFP transgenes were expressed in C4da neurons with the *ppk*-Gal4 driver. VNCs (indicated by the dashed red box in the top panel), which contained transgenic Dscam::GFP in C4da axon terminals, were dissected out from 3^rd^ instar larvae for Western blotting. The bottom panel shows the Western blots of transgenic Dscam::GFP in the axon terminals of C4da neurons. The VNC lysates were used for Western blotting with anti-GFP and anti-Elav antibodies. The relative expression levels are GFP signals normalized by Elav signals. Each dot represents the result from one independent experiment. Overexpression of a [TM1] transgene resulted in a modest level of [TM1] in axon terminals, while two different [TM2] transgenes caused higher levels of [TM2] in axon terminals. Two copies of *ppk*-Gal4 increased the levels of [TM1] in axon terminals. (**H**) The rescue of the axonal defect caused by loss of *Dscam* is proportional to the level of transgenic Dscam in C4da axon terminals, regardless of the isoform. The relative Dscam expression levels were determined by Western blotting of CNS lysates from larvae overexpressing [TM1]::GFP or [TM2]::GFP by *ppk*-Gal4 and plotted against the rescue effect of the transgene.

**Figure 6.**
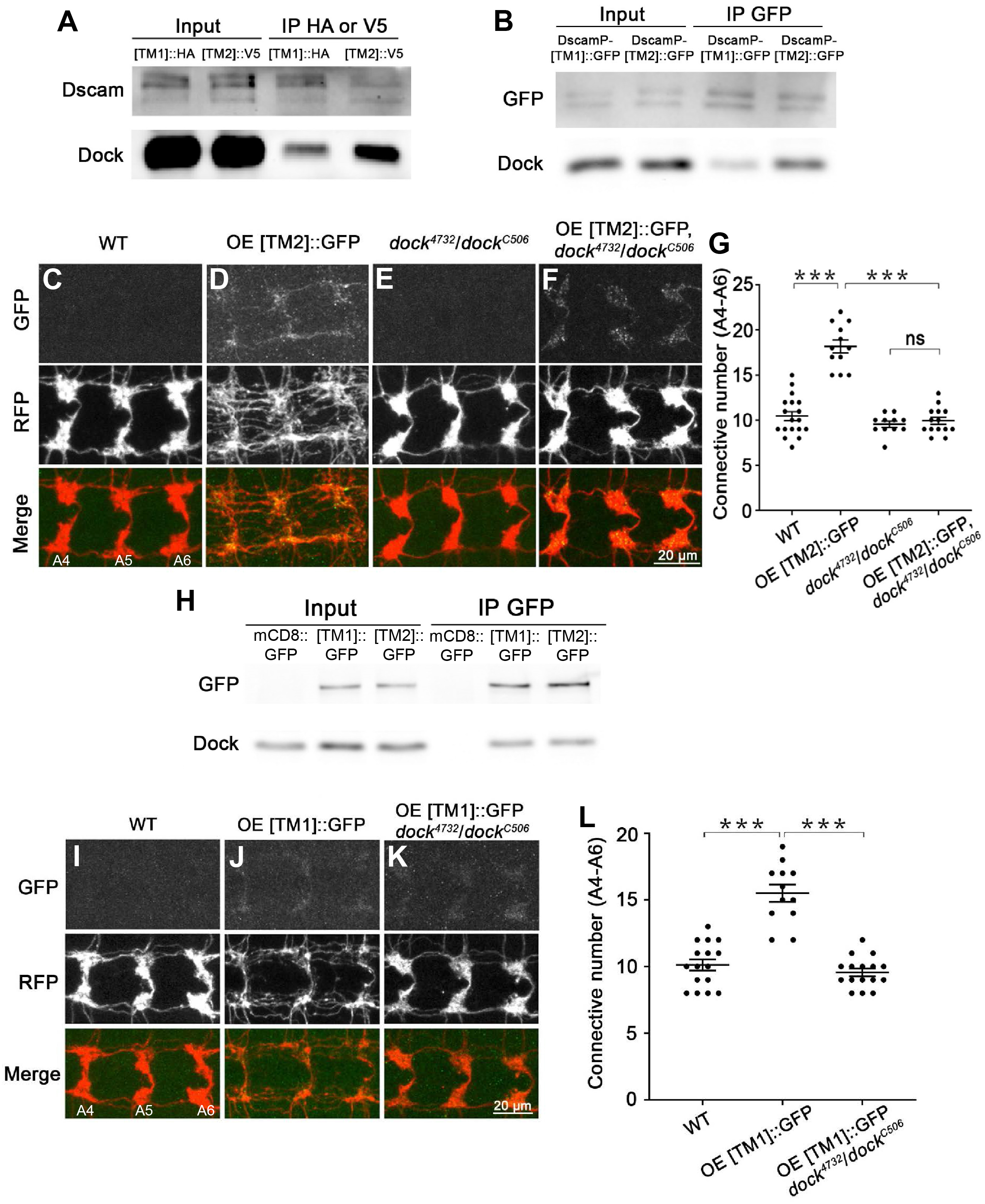
Dscam[TM1] and [TM2] exhibit similar biochemical properties and cellular functions when they are localized in the same subcellular compartment. (**A** & **B**) Dock preferentially interacts with endogenous Dscam[TM2] *in vivo*. (**A**) Brain lysates of larvae with global iso-Tagging of [TM1]::HA or [TM2]::V5 were immunoprecipitated by anti-HA or V5 beads, respectively, and immunoblotted with an anti-Dscam antibody that recognizes both [TM1] and [TM2] and an anti-Dock antibody. This experiment was repeated twice independently. (**B**) Brain lysates of larvae that expressed [TM1]::GFP or [TM2]::GFP through the endogenous *Dscam* promoter were immunoprecipitated by an anti-GFP antibody. The immunoprecipitates were immunoblotted with anti-GFP and anti-Dock antibodies. This experiment was repeated three times independently. (**C-F**) Dscam[TM2] requires Dock to promote presynaptic terminal growth. Shown are representative images of A4-A6. Overexpressing a Dscam[TM2]::GFP transgene significantly promotes axonal growth in C4da neurons (**C** & **D**). While loss of *dock* does not affect C4da axon terminals (**E**), it completely abolishes the overgrowth caused by Dscam[TM2]::GFP overexpression (**F**). (**G**) Quantification of the number of C4da axon connectives. (**H**) Dock binds to [TM1] and [TM2] in similar affinity in cultured S2 cells. Lysates of S2 cells expressing mCD8::GFP, Dscam[TM1]::GFP or [TM2]::GFP were immunoprecipitated with an anti-GFP antibody. Inputs and immunoprecipitates were blotted by anti-GFP and anti-Dock antibodies. This experiment was repeated three times independently. (**I-K**) Transgenic Dscam[TM1] requires Dock to promote presynaptic terminal growth. Compared to WT (**I**), overexpressing [TM1]::GFP transgene significantly promotes axonal growth in C4da neurons (**J**), which is completely abolished by loss of *dock* (**K**). (**L**) Quantification of the number of C4da axon connectives from A4-A6.

Notably, endogenous Dscam expression seemed to be enriched in developing neurons, but not in mature ones. While both endogenous Dscam[TM1] and [TM2] were observed in PNS neurons at the early 2^nd^-instar stage, no signal was detectable in the 3^rd^ instar stage (data not shown). In larval MB, endogenous Dscam[TM2] was detected only in the core fiber, which is the axonal projections of nascent developing neurons. These observations are consistent with previous immunostaining results with an anti-Dscam antibody (Zhan et al., 2004). The discrepancies of spatiotemporal expression pattern between transgenes and endogenous Dscam isoforms underscore the importance of studying protein isoforms at the physiological level.

### Dendrite-specific localization restrains endogenous Dscam[TM1] from functioning in axons

The studies above show that endogenous Dscam[TM1] is required for dendritic, but not axonal, development. Although this could be a result of the compartmentalized localization of Dscam[TM1] in dendrites, trans-compartmental communication has been shown for many membrane proteins (Terenzio et al., 2017). It is thus essential to determine whether or not the dendrite-specific localization of Dscam[TM1], instead of its biochemical properties, restrains it from functioning in axons. This important issue has not been addressed, despite previous insightful studies on the differences between Dscam[TM1] and [TM2].

We determined whether forced localization of Dscam[TM1] in axon terminals was sufficient to rescue the axon phenotype caused by *Dscam[TM2]^iso-KO^*. Overexpression of a *Dscam[TM1]* transgene in C4da neurons led to a modest level of Dscam[TM1] in axon terminals and significantly mitigated axonal growth defects caused by *Dscam[TM2]^iso-KO^* (Figures 5A-B, and F). As expected, two different [TM2] transgenes (#1 and #2) both caused higher levels of [TM2] in axon terminals and stronger rescue than the [TM1] transgene (Figures 5C, E-G). Strikingly, when we used 2 copies of the C4da-specific driver *ppk*-Gal4 to increase the axonal level of Dscam[TM1] to that expressed by *Dscam[TM2]* transgenes, the rescue effects were comparable (Figures 5C, D, F, and G). By quantitatively assessing transgenic Dscam::GFP levels in C4da axon terminals (Figure 5G), we found a linear correlation (R^2^=0.969) between presynaptic Dscam isoform levels and their rescue effects (Figure 5H). These results suggest that when localized in the same subcellular compartment, Dscam[TM1] function equally as Dscam[TM2] in promoting axonal growth. Thus, the dendritic function of endogenous Dscam[TM1] is a result of its compartmentalized localization.

### Axonally enriched Dscam[TM2]-Dock signaling is essential for the axonal function of Dscam[TM2]

While the study above demonstrates that the dendrite-specific localization of Dscam[TM1] prevents it from functioning in axon development, it remains unknown whether Dscam[TM1] and [TM2] exhibit any biochemical difference that might also explain the difference in their cellular functions. To test this possibility, we took advantage of *isoTarget* to compare Dscam[TM1] and [TM2] for their interactions with functional partners. Among the molecules known to bind to the intracellular domain of Dscam and mediate its signaling (Kamiyama et al., 2015; Liu et al., 2009; Purohit et al., 2012; Schmucker et al., 2000; Sterne et al., 2015), we found that Dock, an SH2/SH3 adapter protein that is preferentially localized in axons (Desai et al., 1999; Fan et al., 2003; Hing et al., 1999), preferentially associated with Dscam[TM2] in *Drosophila* larval brains (Figure 6A). We further confirmed this finding with transgenic flies that overexpress Dscam[TM1]::GFP or [TM2]::GFP through their endogenous promoters (Wang et al., 2004). Immunoprecipitation was performed using an anti-GFP antibody. Endogenous Dock preferentially co-precipitated with [TM2]::GFP from brain lysates of 3^rd^ instar larvae (Figure 6B). Consistent with this notion, we found that Dock is required for Dscam[TM2] to promote axon terminal growth. While overexpressing [TM2] significantly promoted axon terminal growth in C4da neurons, its effect was completely eliminated by loss of *dock* (Figures 6C-G). Interestingly, we found that the [TM2]-Dock signaling is compartment-dependent. Dock bound to both isoforms with equal affinity in cultured Schneider 2 (S2) cells, which do not have dendrite-axon compartmentalization (Figure 6H). This result is consistent with the previous finding that the Dock-binding sites, including PXXP sites and polyproline motifs, are located in the cytoplasmic region shared by [TM1] and [TM2] (Schmucker et al., 2000). Consistent with the biochemical results, the axon terminal overgrowth resulted from [TM1] overexpression was also eliminated by loss of *dock* (Figures 6I-L). Thus, given the same subcellular compartment, Dscam[TM1] and [TM2] are biochemically comparable and share the same signaling mechanism. Moreover, the ectopic localization, function and signaling of transgenic [TM1] in axons underscore the importance of studying splicing isoforms at physiological levels of expression.

### Axonal enrichment of Wnd compartmentalizes the Wnd-Dscam[TM2] signaling

We showed previously that the signaling pathway involving the E3 ubiquitin ligase Highwire (Hiw) and the dual leucine zipper kinase Wallenda (Wnd) promotes Dscam expression (Kim et al., 2013). However, whether the Hiw-Wnd pathway regulates the expression of both [TM1] and [TM2] isoforms is unknown. To address this, we determined whether loss of *hiw* or overexpression of Wnd changed the levels of [TM1]::HA and [TM2]::V5. To test the effects of loss of *hiw*, Western blotting was performed on CNS lysates from trans-heterozygous [TM1]::HA / [TM2]::V5 larvae that were generated through global iso-Tagging. We found that loss of *hiw* led to increased expression of both Dscam[TM1] and [TM2] in the CNS (Figure 7A). Next, we compared Wnd’s effect on the expression of Dscam[TM1] and [TM2]. As global overexpression of Wnd caused larval lethality, we co-expressed Wnd and R recombinase (for iso-Tagging of endogenous Dscam[TM1] and [TM2]) in a subset of CNS neurons with GAL4^4-77^ (Kim et al., 2013). Interestingly, Wnd overexpression only increased the level of Dscam[TM2] (Figure 7B), but not that of [TM1], suggesting that unlike Hiw, Wnd preferentially regulates the expression of Dscam[TM2]. This finding from biochemical studies was confirmed in C4da neurons by immunostaining of the endogenous Dscam[TM2] tagged through iso-Tagging. Immunostaining of Dscam[TM2]::V5^iso-Tagging^ showed that endogenous Dscam[TM2] was undetectable in C4da axon terminals in 3^rd^-instar larvae, but detectable at this developmental stage when Wnd was overexpressed in these neurons (Figure 7C).

**Figure 7.**
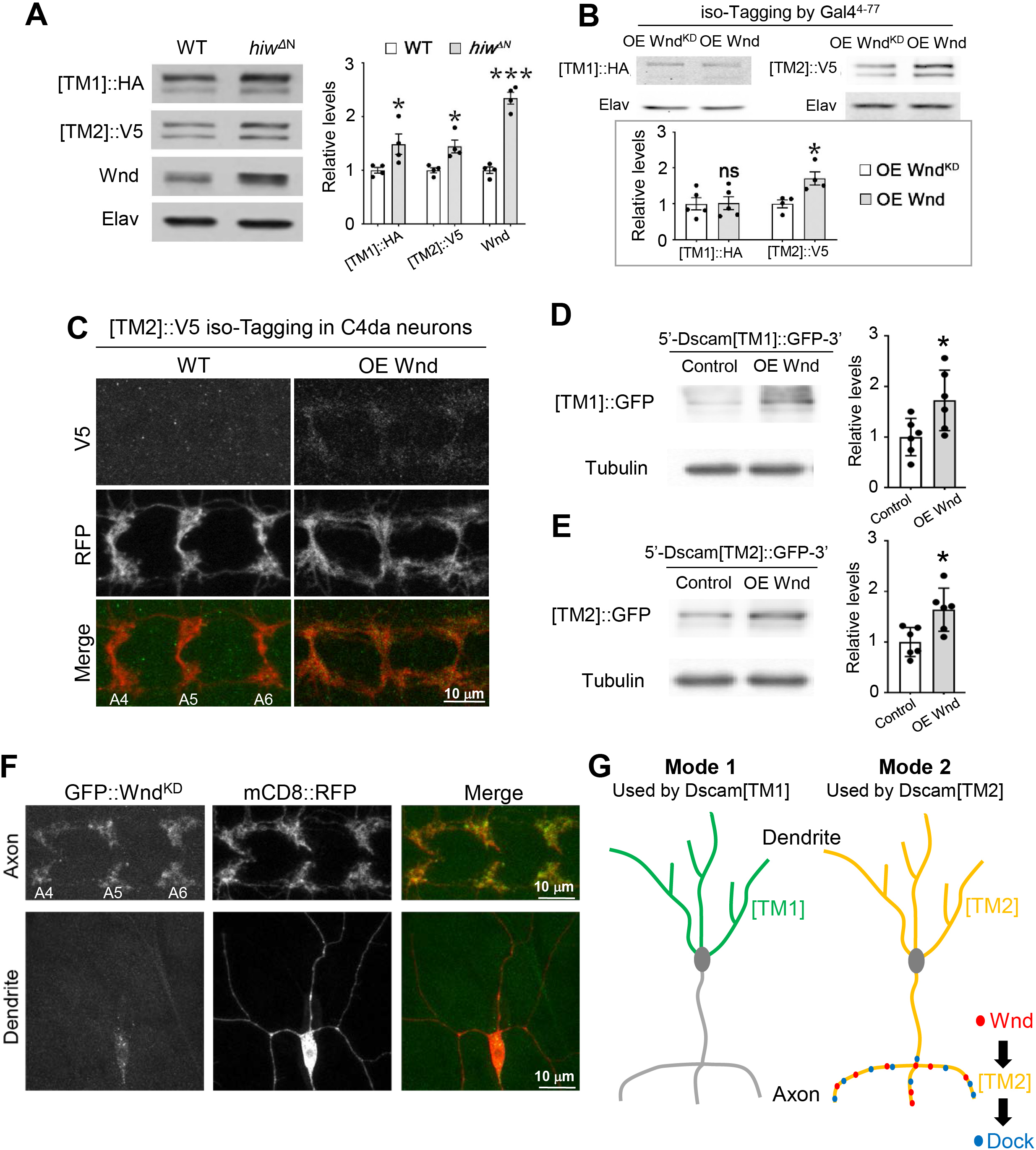
Axonal enrichment of Wnd compartmentalizes Wnd-Dscam[TM2] signaling. (**A**) Hiw suppresses the expression of both Dscam[TM1] and [TM2]. In the brains of 3^rd^ instar larvae that were trans-heterozygotes of global iso-Tagging of [TM1]::HA and [TM2]::V5, loss of *hiw* (*hiw^ΔN^*) elevated the levels of both endogenous [TM1] and [TM2]. Left: Western blots; Right: quantification of Western blots. Each dot represents the result from one independent experiment. (**B**) Overexpression of Wnd increases the levels of Dscam[TM2], but not that of [TM1]. Top: The Gal4^4-77^, which is expressed in a small set of PNS neurons and a number of CNS neurons, was used to drive the expression of R recombinase for iso-Tagging and the overexpression of Wnd. Compared with a kinase-dead Wnd transgene (Wnd^KD^), overexpressing Wnd does not affect endogenous Dscam[TM1] levels (left), but significantly increases Dscam[TM2] levels (right) in 3^rd^-instar larval brains. Bottom: quantification of Western blots. Each dot represents the result from one independent experiment. (**C**) Overexpression of Wnd increases endogenous Dscam[TM2] levels in the presynaptic terminals of C4da neurons. At the 3^rd^ instar larval stage, Dscam[TM2]::V5^iso-Tagging^ is no longer detectable in C4da axon terminals. Overexpression of Wnd elevates the level of Dscam[TM2]::V5^iso-Tagging^ in these terminals. (**D, E**) Wnd similarly promotes [TM1] and [TM2] expression in cultured S2 cells. S2 cells are transfected with plasmids expressing Wnd and [TM1]::GFP or [TM2]::GFP with endogenous *Dscam* 5’ and 3’ UTR. Lysates of S2 cells were blotted by anti-GFP and anti-tublin antibodies. Each dot represents the result from one independent experiment. (**F**) Wnd is enriched in axons. GFP-tagged kinase-dead Wnd (GFP::Wnd^KD^) was expressed in C4da neurons by *ppk*-Gal4. mCD8::RFP was used to label the neurons. While GFP::Wnd^KD^ signal is enriched in axon terminals, little is observed in dendrites. Yellow arrows point to major dendritic branches. (**G**) Subcellular localization expands the functional diversity of splicing isoforms via two different modes. In Mode 1, the localization of an isoform in a particular subcellular location (e.g., Dscam[TM1] in dendrites) restrains this isoform from functioning in other compartments. In Mode 2, the enrichment of the functional partners (e.g., Wnd and Dock) for a ubiquitously localized isoform (e.g., Dscam[TM2]) leads to isoform-specific subcellular signaling and functions.

Next, we investigated the mechanism underlying the differential effect of Wnd on Dscam[TM1] and [TM2]. Firstly, we determine whether Wnd is capable of increasing Dscam[TM1] expression in a cell type without the dendrite-axon compartmentalization. We overexpressed Dscam isoforms with endogenous 5’ and 3’ UTR in S2 cells, and found that Wnd similarly promoted [TM1] and [TM2] expression (Figure 7D-E). This result suggest that Dscam[TM1] and [TM2] are molecularly indistinguishable for the Wnd regulation. Secondly, we identified the distribution of Wnd in neurons, and found that Wnd was enriched in axons but not dendrites. A GFP-tagged kinase-dead version of Wnd (GFP-Wnd^KD^)(Xiong et al., 2010) was expressed in C4da neurons; substantial signals were detected in axon terminals but not in dendrites (Figure 7F). These data suggest that axonal localization of Wnd compartmentalizes the Wnd-Dscam[TM2] signaling.

## DISCUSSION

In this study, we developed *isoTarget* to generate splicing isoform-specific loss-of-function mutants and conditional tagging in specific neurons. As a proof of concept, we applied *isoTarget* to investigate two mutually exclusive isoforms of Dscam, which differ in their transmembrane and juxtamembrane domains, for their subcellular localization, regulation of expression, function in dendrite/axon development, and subcellular signaling. These findings highlight the versatility of *isoTarget* and the importance of studying splicing isoform at endogenous levels *in vivo*. In addition, they establish the causality between the subcellular localization and cellular function of splicing isoforms, demonstrating the critical role of subcellular localization in expanding the functional diversity of splicing isoforms.

Our study demonstrate that Dscam isoforms use two different modes to achieve distinct compartment-specific functions *in vivo* (Figure 7G). While the dendrite-specific localization restrains Dscam[TM1] from functioning in axons, axonal enrichment of functional partners forms a subcellular signaling pathway involving the ubiquitously distributed Dscam[TM2].

### Subcellular localization defines the cellular functions of alternative splicing isoforms that have the same biochemical functions

Alternative isoforms often differ in their protein structures and thus the biochemical properties (Kelemen et al., 2013). Recent studies suggest the importance of subcellular localization in defining the cellular functions of alternative splicing isoforms. For example, in cultured neurons, the RBFOX1 gene generates a nuclear variant (by excluding exon 19) that regulates RNA splicing and a cytoplasmic variant (by including exon 19) that stabilizes mRNAs (Lee et al., 2016). However, whether subcellular localization is the only reason why the nuclear and cytoplasmic variants differ in their cellular functions remains to be determined, as it is possible that the two isoforms have distinct biochemical functions. More broadly, a challenging question in the field is whether the difference in subcellular localization determines the compartmentalized functions for the isoform (i.e., the issue of causality), especially at the endogenous levels *in vivo*. By applying *isoTarget* in *Drosophila* sensory system, we establish the causality between the subcellular localization and cellular functions of the Dscam isoforms. Our findings highlight the critical role of subcellular localization in expanding the functional diversity of isoforms.

Even if they are localized in different subcellular compartments, splicing isoforms with the same biochemical function might not exhibit distinct cellular functions because their downstream partners might spread throughout the cell (e.g., via trans-compartmental communication (Terenzio et al., 2017)). In order to achieve cellular functions specific to a compartment, one solution is to localize the functional partners shared by the isoforms to specific subcellular compartments. Indeed, we found that compartmental enrichment of interactors leads to isoform-specific signaling. The Dscam[TM1] and [TM2] do not differ in their biochemical interactions with Wnd and Dock (Figure 6H, 7C-D). Yet, the axonal enrichment of Wnd and Dock forms a compartmentalized Wnd-Dscam[TM2]-Dock signaling cascade *in vivo*, despite that Dscam[TM2] is present in both dendrites and axons (Figure 4C, E, H, K). These findings suggest that, for splicing isoforms with the same biochemical functions, specific cellular functions can be achieved by the compartment-specific colocalization of the isoforms and their functional partners (Figure 7G).

Sphingolipids are essential for trafficking of dendritic and axonal cell adhesion molecules, including Dscam (Goyal et al., 2019). Loss of SPT, a key enzyme in sphingolipid biosynthesis, reduces the membrane localization of Dscam[TM1] and [TM2] in subsets of mushroom body neurons, resulting in [TM1] aggregates in the soma and [TM2] aggregates in the axons. *isoTarget* would be instrumental for testing the function of sphingolipids in the trafficking of endogenous Dscam isoform in select neuron populations. Moreover, the *isoTarget-based* approaches described in our paper allowed us to address whether subcellular localization is the cause of the isoform-specific function. These approaches can also be used to determine the causality between alternations in subcellular localization (e.g., those caused by trafficking defects) and changes in cellular functions.

### Advantages of *isoTarget*

The studies reported in this paper demonstrate several advantages of *isoTarget* over traditional techniques.

First, it can be used to generate classic genetic mutants for analyzing the functions of specific splicing isoforms. RNA interference has been adopted to investigate Dscam isoform functions in CNS neurons (Shi et al., 2007). However, we found that this method is not applicable for studying Dscam in larval PNS neurons (data not shown). This is likely because endogenous Dscam primarily functions at early developmental stages and a late efficacy of RNAi precluded the discovery of Dscam functions in these neurons. By contrast, *isoTarget* can be used to create a loss-of-function mutant a specific isoform without affecting other isoforms, allowing us to investigate the functions of Dscam isoforms in larval PNS neurons.

Second, *isoTaget* enables the identification of isoform-specific localization at the subcellular level in neurons of interest *in vivo*, which is otherwise challenging due to the difficulty in discerning immunofluorescent signals in subcellular compartments among a number of neurons expressing the same protein in the vicinity. In addition, this method expresses tagged proteins at more physiological levels than transgenes. Prior studies on Dscam have relied on transgenes to investigate the functions and subcellular localizations of Dscam[TM1] and [TM2] (Goyal et al., 2019; Kim et al., 2013; Soba et al., 2007; Wang et al., 2004; Yang et al., 2008; Zhan et al., 2004), but transgenic proteins are often ectopically localized. For example, whereas the endogenous Dscam[TM1] is absent in axons (Figures 4D and S5B), overexpressing [TM1] leads to its axonal localization (Figures 5A-G). In fact, tagging endogenous Dscam[TM1] and [TM2] with *isoTarget* led to the observation of consistent patterns of subcellular localization in different types of neurons (Figure 4), which reconciles the cell-type discrepancies observed previously with transgenes.

Third, *isoTarget* allows studying isoform-specific compartmentalized signaling. Using *isoTarget*, we identified a Wnd-Dscam[TM2]-Dock signaling pathway enriched at the presynaptic terminals of C4da neurons. Complementing the *in vivo* studies with *isoTarget*, we performed biochemical studies in S2 cells and found that Dscam[TM1] and [TM2] did not differ in their biochemical interactions with Wnd and Dock (Figures 6H, 7C-D). Consistent with this, forced localization of ectopic Dscam[TM1] in axons also increased axonal growth *in vivo* (Figures 5A-G) through the same downstream effector Dock used by Dscam[TM2] (Figures 6I-L). In this series of studies, *isoTarget* was essential for establishing the cellular functions and biochemical interactions *in vivo*.

### Limitations

There are three limitations of the *isoTarget* technique. First, for the successful application of *isoTarget*, the isoform inserted with iso-KO cassette is expected to lose its function. This is not necessarily always the case, especially when the targeted exon encodes a fragment located at the C-terminus of the protein. This problem is common in isoform studies by genetic modifications, including Cre-LoxP and isoEXPRESS (Gu et al., 2019). Developing isoform-specific nanobody might be a way to solve this problem (Roth et al., 2019). Second, we discovered that the original translational stop cassette may cause off-target effects (Figures S3A-C and F), and that such effects depend on the length of the cassette. Thus, the expression of isoforms other than the targeted one should always be examined by techniques such as RT-qPCR. Finally, successful uses of *isoTarget* require that the epitope tagging preserves the function of the splicing isoform. Structural information would be helpful in choosing the proper inserting site. For example, inserting the isoTarget cassette into the loop region of a polypeptide is likely to increase the chance of success.

In summary, we have developed *isoTarget* as a versatile genetic tool that is compatible with a variety of techniques for analyzing isoform-specific properties and uncovering the mechanisms underlying isoform diversity at multiple levels *in vivo*. We anticipate this methodology be useful for isoform studies in various cell types and organisms.

## ACKNOWLEDGEMENT

We thank Drs. Kenneth Kwan, Ken Inoki, Yukiko Yamashita, Dawen Cai for helpful discussions, Dr. Jung Hwan Kim for teaching H.L. and for critical suggestions on *isoTarget* validation. We also thank Drs. Tzumin Lee, Larry Zipursky, Yi Chen, Jack Dixon, Catherine Collins, Ryan Insolera, Scott Barolo, and David Lorberbaum for sharing reagents. We thank Drs. Catherine Collins, Laura Smithson, and Elizabeth Cebul for their critiques on an earlier version of the manuscript. This work was supported by grants from NIH (R01 MH092147 and R01 NS095525 to BY) and Protein Folding Disease Initiative of the University of Michigan to B.Y. M.W.Z. was supported by the NIH Cellular and Molecular Biology Training Grant T32-GM007315.

## AUTHOR CONTRIBUTIONS

H.L. and B.Y. conceived the project and designed the experiments. H.L. designed, generated and validated *isoTarget* flies, examined the functions, endogenous expression and signaling cascade of Dscam isoforms. S.P., R.L. and M.W.Z. performed MARCM on C4da neurons and assisted in quantification of global iso-KO. W.Z. examined isoform functions in C3da neurons. Y. H. assisted in generating *isoTarget* flies. L.Y. assisted in experiments of endogenous isoform expression. B.Y. supervised the project. H.L. and B.Y. wrote the paper.

## EXPERIMENTAL PROCEDURES

### *Drosophila* strains

The following published fly strains were used: *ppk*-Gal4 (chromosome III) (Kuo et al., 2005); GMR83B04-GAL4 (Pfeiffer et al., 2011); UAS-Dscam[TM2]::GFP (3.36.25, chromosome III and X), UAS-Dscam[TM1]::GFP (3.36.25, chromosome X), *DscamP*-Dscam[TM1]::GFP (3.36.25) (Wang et al., 2004), *DscamP*-Dscam[TM2]::GFP (3.36.25) (Wang et al., 2004); OK107-Gal4 (Connolly et al., 1996); *Dscam^18^* (Wang et al., 2002); *hiw^ΔN^* (Wu et al., 2005); *wnd^1^, wnd^3^*, and UAS-Wnd (Collins et al., 2006); *dock^4732^* and *dock^C506^* (Garrity et al., 1996); *nos*-Gal4 (Van Doren et al., 1998); 20xUAS-R recombinase (Nern et al., 2011).

### Generation of DNA constructs

The design of *isoTarget* system is described in Figure S1A. The translational stop cassette was amplified by PCR from non-catalytic region of lacZ with frame shift to introduce multiple premature termination codons. The loxP-dsRed-loxP sequence is described in Gratz et al., (2014) (Gratz et al., 2014). The pBluescript donor plasmid, which contains *isoTarget* KI cassettes (GS linker-RSRT-*tlstop*-loxP-dsRed-loxP-RSRT-epitope-G3 linker) and Dscam isoform homologous sequences with mutated PAM, was generated with the In-Fusion HD Cloning Kit (Clontech Laboratories, Inc.). The pCFD3-dU63gRNA plasmid, which produces gRNA in fly embryos, is described in Ran et al., (2013) (Ran et al., 2013). The pUAST-*Dscamδ’UTR-Dscam[TM1]::GFP-Dscam3’UTR* (4.3-6.36-9.25) plasmid was made by modifying pUAST-*Dscamδ’UTR-Dscam[TM2]::GFP-Dscam3’UTR* (4.3-6.36-9.25)(Kim et al., 2013). Specifically, the fragment containing *Dscam*5’UTR and exons 1-16 and that containing exons 18-24 plus *Dscam3’UTR* were amplified from pUAST-*Dscamδ’UTR-Dscam[TM2]::GFP-Dscam3’UTR* by PCR. The fragments, together with PCR products of *Dscam* exon 17.1, were fused with the pUASTattB vector (linearized by EcoRV) by In-Fusion HD Cloning (Clontech Laboratories, Inc.).

### Sequences of *isoTarget* cassettes

<u>*tlstop*</u> (in-frame stop codons that we introduced are underlined): <u>TAA</u>CGTAAGCTAGCTAGACCGGTCCCAACT<u>TAA</u>TCGCCTTGCAGCACATCCCCCTT TCGCCAGCTGGCG<u>TAATAG</u>CGAAGAGGCCCGCACCGATCGCCCTTCCCAACAGTT GCGCAGCCTGAATGGCGAATGGCGCTTTGCCTGGTTTCCGGCACCAGAAGCGGT GCCGGAAAGCTGGCTGGAGTGCGATCTTCC<u>TGA</u>GGCCGATACTGTCGTCGTCCCC TCAAACTGGCAGATGCACGGTTACGATGCGCCCATCTACACCAACGTAACCTATCC CATTACGGTCAATCCGCCGTTTGTTCCCACGGAGAATCCGACGGGTTGTTACTCGC TCACATT<u>TAA</u>TGT<u>TGATGA</u>AAGCTGGCTACAGGAAGGCCACGCGTA

<u>Short *tlstop*</u> (in-frame stop codons that we introduced are underlined): <u>TAA</u>CGTAAGCTAGCTAGACCGGTTTCCCACGGAGAATCCGACGGGTTGTTACTCG CTCACATT<u>TAA</u>TGT<u>TGATGA</u>AAGCTGGCTACAGGAAGGCCACGCGTA

<u>GS-linker</u>: GGTGGCGGCGGAAGCGGAGGTGGAGGCTCC

<u>RSRT</u>: CTTGATGAAAGAATAACGTATTCTTTCATCAAG loxP: ATAACTTCGTATAATGTATGCTATACGAAGTTAT

<u>dsRed</u> box (consisting of 3XP3 enhancer/HSP70 promoter (Gratz et al., 2014), the cDNA encoding the <u>dsRed fluorescent protein</u> and *SV40 3’ UTR*) CGTACGGGATCTAATTCAATTAGAGACTAATTCAATTAGAGCTAATTCAATTAGGAT CCAAGCTTATCGATTTCGAACCCTCGACCGCCGGAGTATAAATAGAGGCGCTTCGT CTACGGAGCGACAATTCAATTCAAACAAGCAAAGTGAACACGTCGCTAAGCGAAAG CTAAGCAAATAAACAAGCGCAGCTGAACAAGCTAAACAATCGGCTCGAAGCCGGT CGCCACC<u>ATGGCCTCCTCCGAGGACGTCATCAAGGAGTTCATGCGCTTCAAGGTG CGCATGGAGGGCTCCGTGAACGGCCACGAGTTCGAGATCGAGGGCGAGGGCGA GGGCCGCCCCTACGAGGGCACCCAGACCGCCAAGCTGAAGGTGACCAAGGGCG GCCCCCTGCCCTTCGCCTGGGACATCCTGTCCCCCCAGTTCCAGTACGGCTCCAA GGTGTACGTGAAGCACCCCGCCGACATCCCCGACTACAAGAAGCTGTCCTTCCCC GAGGGCTTCAAGTGGGAGCGCGTGATGAACTTCGAGGACGGCGGCGTGGTGACC GTGACCCAGGACTCCTCCCTcCAGGACGGCTCCTTCATCTACAAGGTGAAGTTCAT CGGCGTGAACTTCCCCTCCGACGGCCCCGTAATGCAGAAGAAGACTATGGGCTG GGAGGCgTCCACCGAGCGCCTGTACCCCCGCGACGGCGTGCTGAAGGGCGAGAT CCACAAGGCCCTGAAGCTGAAGGACGGCGGCCACTACCTGGTGGAGTTCAAGTC CATCTACATGGCCAAGAAGCCCGTGCAGCTGCCCGGCTACTACTACGTGGACTCC AAGCTGGACATCACCTCCCACAACGAGGACTACACCATCGTGGAGCAGTACGAGC GCGCCGAGGGCCGCCACCACCTGTTCCTGTAG</u>*GGGCCGCGACTCTAGATCATAAT CAGCCATACCACATTTGTAGAGGTTTTACTTGCTTTAAAAAACCTCCCACACCTCCCCCTGAACCTGAAACATAAAATGAATGCAATTGTTGTTGTTAACTTGTTTATTGCAGCTTATAATGGTTACAAATAAAGCAATAGCATCACAAATTTCACAAATAAAGCATTTTTTTCACTGCATTCTAGTTGTGGTTTGTCCAAACTCATCAATGTATCTTAACCGGT*

<u>V5</u>: GGCAAGCCCATCCCAAACCCACTGCTCGGCCTGGATAGCACC

<u>HA</u>: TACCCATACGATGTTCCAGATTACGCT

<u>G3 linker</u> (Gratz et al., 2014): GGTGGCGGC

### Generation of *isoTarget* flies

The donor plasmid (750 ng/μl) and gRNA plasmid (250 ng/μl) were co-injected into fly embryos to generate mosaic G0 flies, which were crossed to *white*^-/-^ flies to get G1 heterozygous knock-in flies that express the selection marker dsRed in their eyes. G1 flies with red fluorescence in their eyes were crossed to Cre-expressing flies (BL1092) to generate iso-KO flies. To generate global iso-Tagging flies, iso-KO flies were mated with the germline driver *nos*-Gal4 (BL4442) and 20x UAS-R recombinase (BL55795). According to our experiences, around 20% of fertile mosaic G0 flies were able to generate G1 knock-in flies with fluorescent red eyes. The G1 knock-in flies constitute 1-30% of total G1 flies, depending on individual lines. To generate cell-type-specific iso-Tagging, iso-KO flies were mated with cell-type-specific Gal4 drivers, such as *ppk*-Gal4, and 20x UAS-R recombinase.

### S2 cell culture and transfection

S2 cells were maintained in *Drosophila* Schneider’s medium supplemented with 10% fetal bovine serum at 25°C in a humidified chamber. Plasmids were transfected into cultured S2 cells with polyethylenimine (PEI) (Ehrhardt et al., 2006). Cells were harvested for Western blotting 2 days after transfection.

### Co-immunoprecipitation and Western blotting

To perform co-immunoprecipitation with neural tissues, the CNS of 3^rd^-instar larvae (~120 per experimental condition) were dissected out and placed in ice-cold PBS containing 2 mM sodium vanadate. After a pulse of centrifugation, larval CNS were isolated and then lysed on ice with the lysis buffer (50 mM Tris-HCl/pH 7.4, 150 mM NaCl, 2 mM sodium vanadate, 10 mM sodium fluoride, 1% Triton X-100, 10% glycerol, 10 mM imidazole and 0.5 mM phenylmethylsulfonyl fluoride). Lysates were centrifuged at 15,000 g for 20 min at 4°C. The resulting supernatants were either saved as inputs or incubated with magnetic beads conjugated with appropriate antibodies for 4 hours at 4°C. After washing once with the lysis buffer, twice with lysis buffer containing 0.1% deoxycholate, and 3 times with lysis buffer lacking Triton X-100, the immunoprecipitates and total lysates were resolved on 7.5% SDS-PAGE gels followed by Western blotting as previously described (Kim et al., 2013). Protein samples were transferred to nitrocellulose membranes and detected by chemiluminescence (Catalog# 32106, Pierce ECL Western Blotting Substrate) with either a BIO-RAD ChemiDoc Touch Imaging system or a Kodak X-OMAT 2000 film processor.

The procedure of co-immunoprecipitation with lysates from S2 cells was similar except that cultured S2 cells were re-suspended in ice-cold PBS before lysis.

### Imaging and image analysis

Larvae immunostaining was described previously (Ye et al., 2011). Confocal imaging was done with a Leica SP5 confocal system with 20x or 63x glycerol immersion lenses. To minimize the variation, we only imaged neuronal dendrites and presynaptic terminals of C3da or C4da neurons in abdominal segments 4, 5, and 6. Images were collected with z stacks of 1-μm-step size for dendrites and 0.3-μm-step size for axons. The resulting three-dimensional images were projected into two-dimensional images by maximum projection. The same imaging setting was applied throughout the imaging process.

The Neurolucida software was used to quantify dendritic morphology and axon terminal growth. For dendrites, the overlap of sister branches is counted as a cross. For quantifying axon terminals of single C4 da neuron, branches shorter than 5 μm were excluded from the analysis. For quantifying the number of longitudinal axonal branches visible between abdominal segment 4-6 (i.e., the connectives) of C4 da neurons, only complete connectives spanning neighboring segments were quantified. Fasciculated connectives are counted as two.

To eliminate experimenter’s bias, these experiments were carried out in double-blind fashion. The images acquired by the primary experimenter were coded and randomized by another lab member. After the primary experimenter quantified the data, the data were decoded for statistical analysis.

### Reverse-transcription real-Time PCR

The procedure was as described before (Kim et al., 2013). Briefly, mRNA was extracted from around 20 3rd-instar larval CNS with a standard Trizol method (Invitrogen). cDNA was synthesized with Invitrogen SuperScript III First-Strand Synthesis SuperMix (Invitrogen). 10 ng cDNA was used as the template for each real-time PCR reaction by SYBR Green mix (Thermo Scientific) with the Applied Biosystems 7300. To normalize *Dscam* transcripts to those of reference genes, we calculated ΔCt (*Dscam*) = Ct (*Dscam*) – Ct (reference gene). Statistical analysis was performed to compare ΔCt (*Dscam*)(wild-type) and ΔCt (*Dscam*)(mutant) to determine whether there is the expression of test gene is different between wild-type and mutants (Yuan et al., 2006). Relative mRNA levels is calculated as: 2^-ΔΔCt^, where ΔΔCt (*Dscam*) = ΔCt (*Dscam*)(mutant) -ΔCt (*Dscam*)(wild-type). We used *elav* as the reference gene and *Chmp1* as the internal control. Following primers were used: *Chmp1:* 5’-AAAGGCCAAGAAGGCGATTC-3’ and 5’-GGGCACTCATCCTGAGGTAGTT-3’; *elav*: 5’-CTGCCAAAGACGATGACC-3’ and 5’-TAAAGCCTACTCCTTTCGTC-3’; *Dscam[TM1]*: 5’-CGTTACCGGAGGCACTATCG-3’ and 5’-ATCGTCTTTGTGGTGA TTGCC-3’; *Dscam[TM2]*: 5’-CGTTACCGGAGGCACCATT-3’ and 5’-ACTACATCG TAGTACACATCCTTT-3’.

### Statistical Analysis

Data are presented as mean ± SEM. Comparisons of mean differences between groups were performed by One-way ANOVA followed by Student’s *t*-test. P < 0.05 was considered to be statistically significant. For all quantification, *: p < 0.05; **: p < 0.01; ***: p < 0.001; ns: not significant (p > 0.05).

