## Supplemental Information for "*isoTarget*: a genetic method for analyzing the functional diversity of splicing isoforms *in vivo*"

### Inventory of Supplemental Information

### Supplemental Data

Figure S1. The design of the *isoTarget* cassette.
related to Figure 1.

Figure S2. Validation of *isoTarget* in Dscam[TM2].
related to Figure 1B and 2A-B.

Figure S3. Validation of *isoTarget* in Dscam[TM1] and the discovery of short tlstop cassette.
related to Figure 1B and 2A-B .

Figure S4. Dscam[TM1] and [TM2] function redundantly in mediating dendritic self-avoidance in C4da neurons.
related to Figure 3.

Figure S5. Endogenous Dscam[TM2], but not [TM1], is localized in axons connecting the CNS and PNS.
related to Figure 4.

### Supplemental Figure Legends

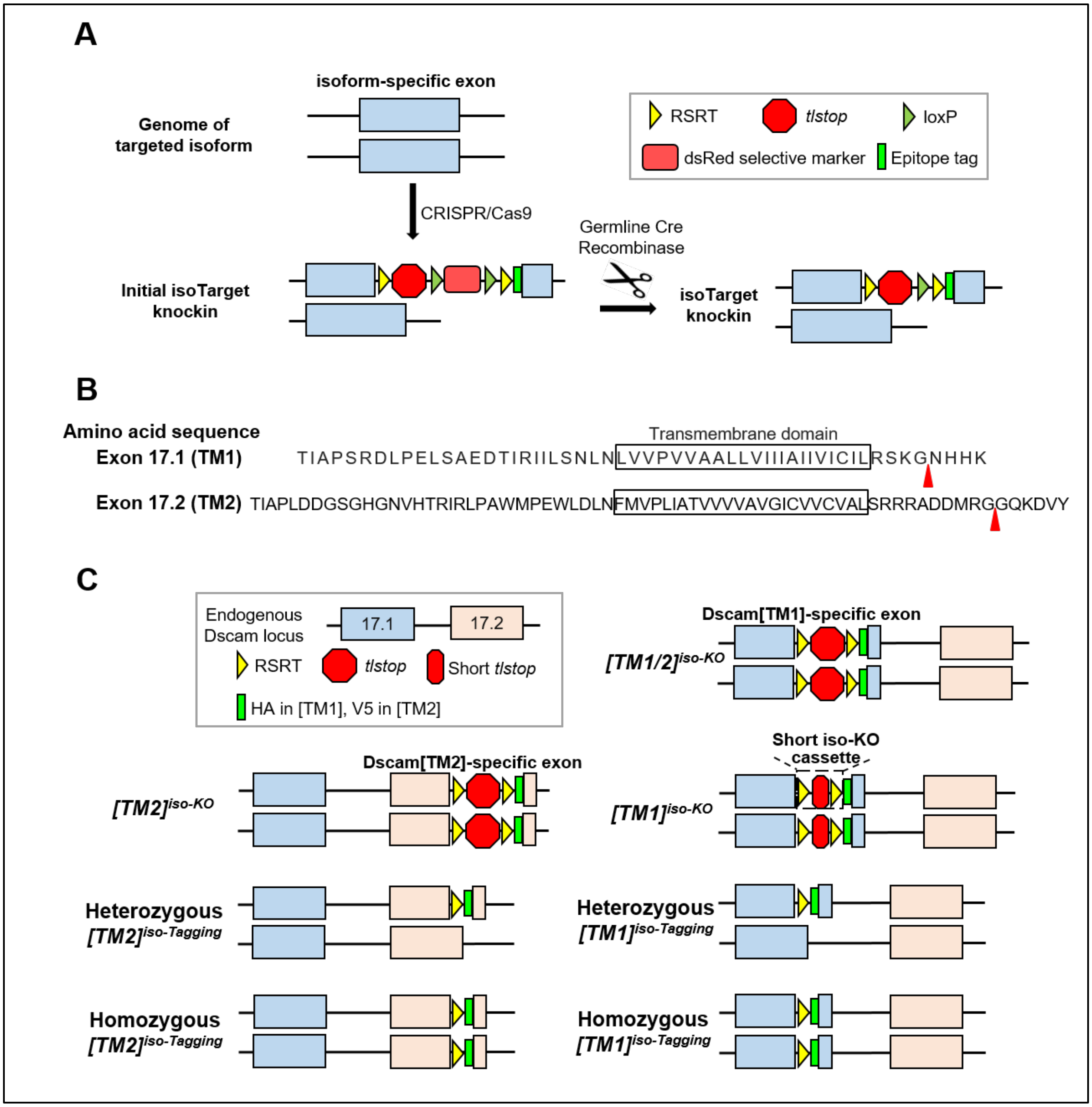


**Figure S1. The design of the *isoTarget* cassette. (Related to Figure 1)**

(**A**) At the core of the cassette is a *tlstop* that contains multiple termination codons to ensure the disruption of mRNA translation. To achieve cell-type-specific labeling of targeted isoforms through inducible site-specific recombination, the *tlstop* is flanked by two RSRT sites followed by an epitope tag. To facilitate the screening of animals with successful integration of the *isoTarget* cassette, a transgene expressing dsRed under the 3xP3 enhancer/Hsp70 promoter is inserted in between the *tlstop* and the second

RSRT site. The dsRed transgene is flanked by two loxP sites, allowing the removing of the dsRed in the presence of Cre recombinase. The *isoTarget* cassette can be knocked into the exon encoding the targeted isoform by CRISPR/Cas9.

(**B**) The *isoTarget* insertion sites in Dscam[TM1] and [TM2] isoforms. Shown are the amino acid sequences encoded by the 17.1 (TM1) and 17.2 (TM2) exons. Red arrow heads indicate the insertion sites in the juxtamembrance regions.

(**C**) Application of *isoTarget* to studying Dscam isoforms. As shown later, iso-KO of [TM1] impairs both [TM1] and [TM2] expression, and is thus named $[TM1/2]^{iso\text{-}KO}$. We discovered a short *tlstop* cassette ("short iso-KO") that specifically knocks out [TM1], but not [TM2]. Iso-Tagging of Dscam[TM1] does not affect [TM1] functions. Iso-KO of [TM2] abolishes, while iso-Tagging preserves, Dscam[TM2] functions. Depending on the purpose of the experiment, Dscam[TM1] and [TM2] can be tagged either globally or specifically in targeted cell types.

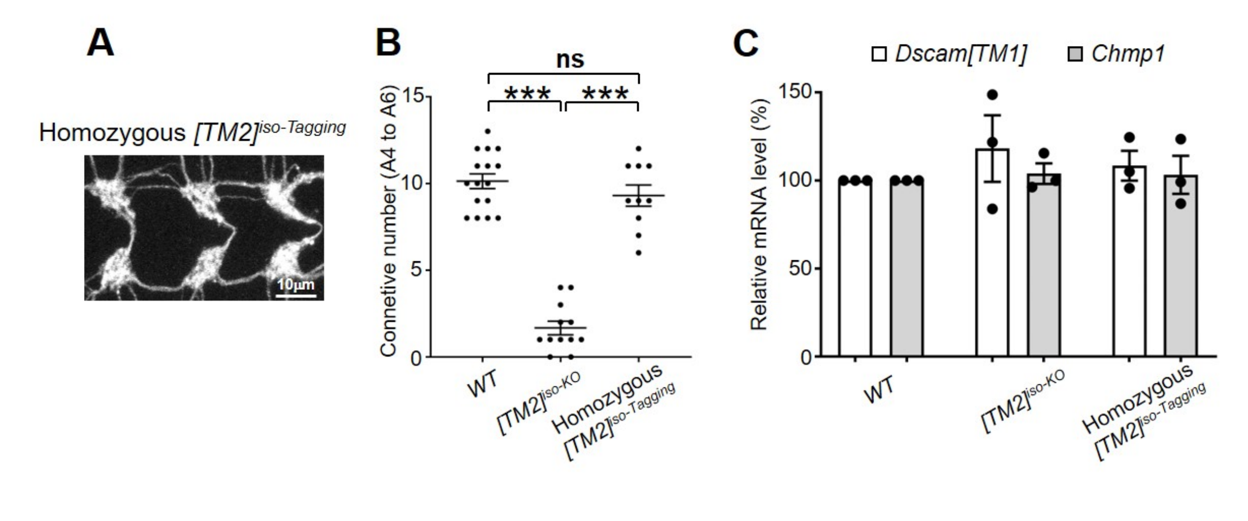


**Figure S2. Validation of *isoTarget* in Dscam[TM2]. (Related to Figure 1B and 2A-B)**

(**A**) *[TM2]*$^{iso\text{-}Tagging}$ does not affect [TM2] functions. As shown in Figure 2A and B, *[TM2]*$^{iso\text{-}KO}$ impairs axon terminal growth in C4da neurons. The axonal defect is completely rescued in homozygous global *[TM2]*$^{iso\text{-}Tagging}$ larvae generated by excision of the RSRT-tlstop-RSRT box from the iso-KO. Shown are the C4da axon terminals in A4-A6 segments.

(**B**) Quantification of the number of C4da axon connectives.

(**C**) Neither global *[TM2]*$^{iso\text{-}KO}$ nor homozygous global *[TM2]*$^{iso\text{-}Tagging}$ affect the mRNA levels of *Dscam[TM1]*. mRNAs from the whole CNS of 3$^{rd}$ instar larvae were used for reverse-transcription real-time PCR. The $\Delta$Ct value of [TM2] mRNA is normalized to that of *elav* gene as descried in the Experimental Procedure. *Chmp1* serves as the internal control. Each dot represents one independent experiment.

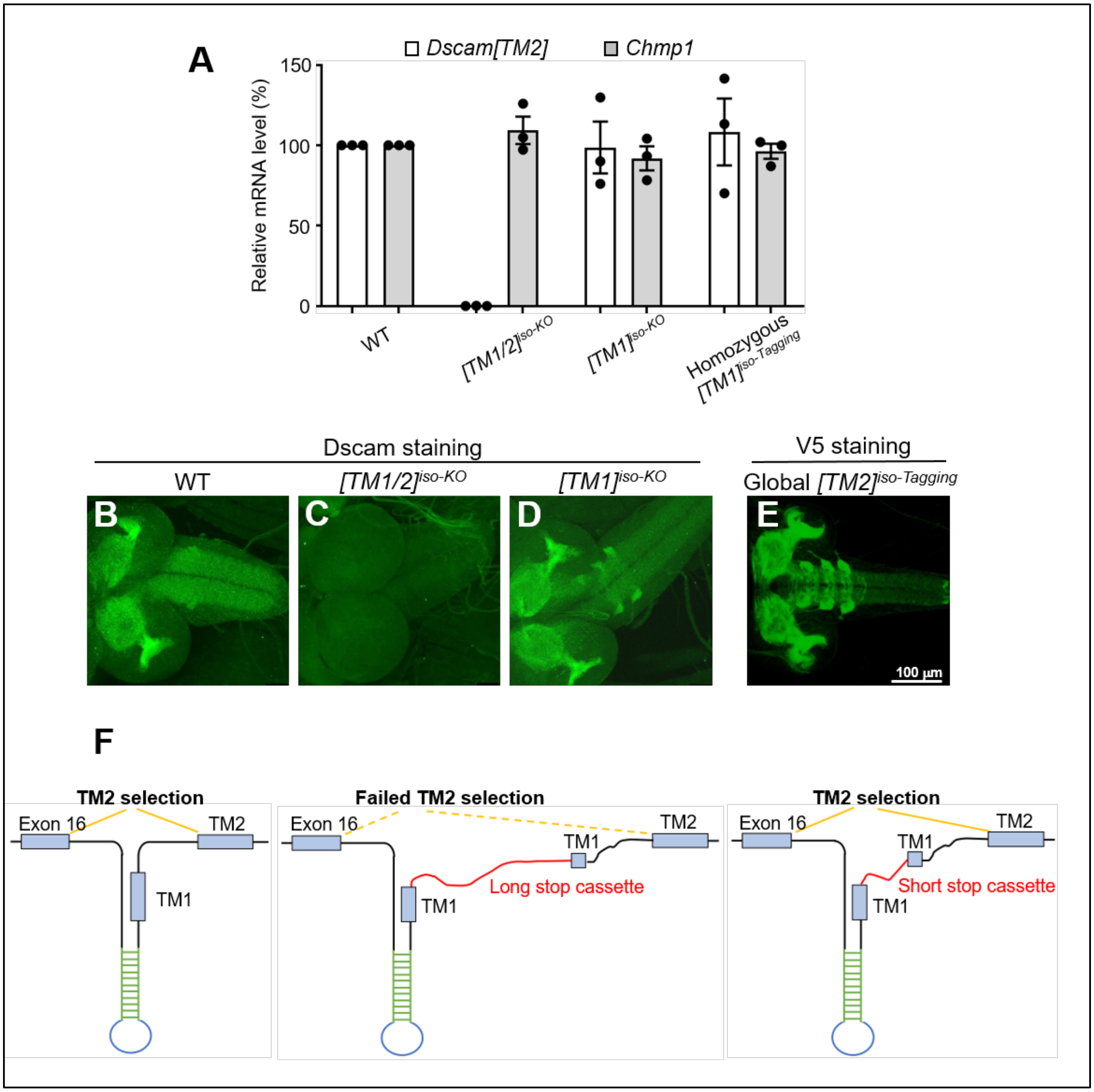


**Figure S3. Validation of *isoTarget* in Dscam[TM1] and the discovery of short *tlstop* cassette. (Related to Figure 1B and 2A-B)**

(**A**) *Dscam[TM2]* mRNA is undetectable in global *Dscam[TM1]* isoTarget flies resulted from the insertion of the original long *tlstop* cassette ("*[TM1/2]*$^{iso\text{-}KO}$"), but remains intact in global *Dscam[TM1] isoTarget* flies generated by the insertion of the short *tlstop* cassette ("*[TM1]*$^{iso\text{-}KO}$"). *Dscam[TM2]* mRNA level is not affected in homozygous global

*[TM1]^{iso-Tagging}* larvae. mRNAs from the whole CNS of 3rd instar larvae were used for reverse-transcription real-time PCR.

(**B**-**E**) Immunostaining of the CNS with an anti-Dscam antibody shows that while Dscam expression is abolished in global *Dscam[TM1/2]^{iso-KO}* (**C**), it is detectable in global *Dscam[TM2]^{iso-KO}* (**D**) and exhibits a pattern that is similar to endogenous Dscam[TM2] labeled by global [TM2]::V5 iso-Tagging (**E**).

(**F**) A schematic model of TM exon selection from *Dscam* pre-mRNA, based on a previous study (Anastassiou et al., 2006). The complementary sequences (green) form a stem-loop, which is required for the selection of TM2 exon into *Dscam* mRNA. Inserting a long cassette into TM1 exon may create a long distance that eliminates TM2 exon selection.

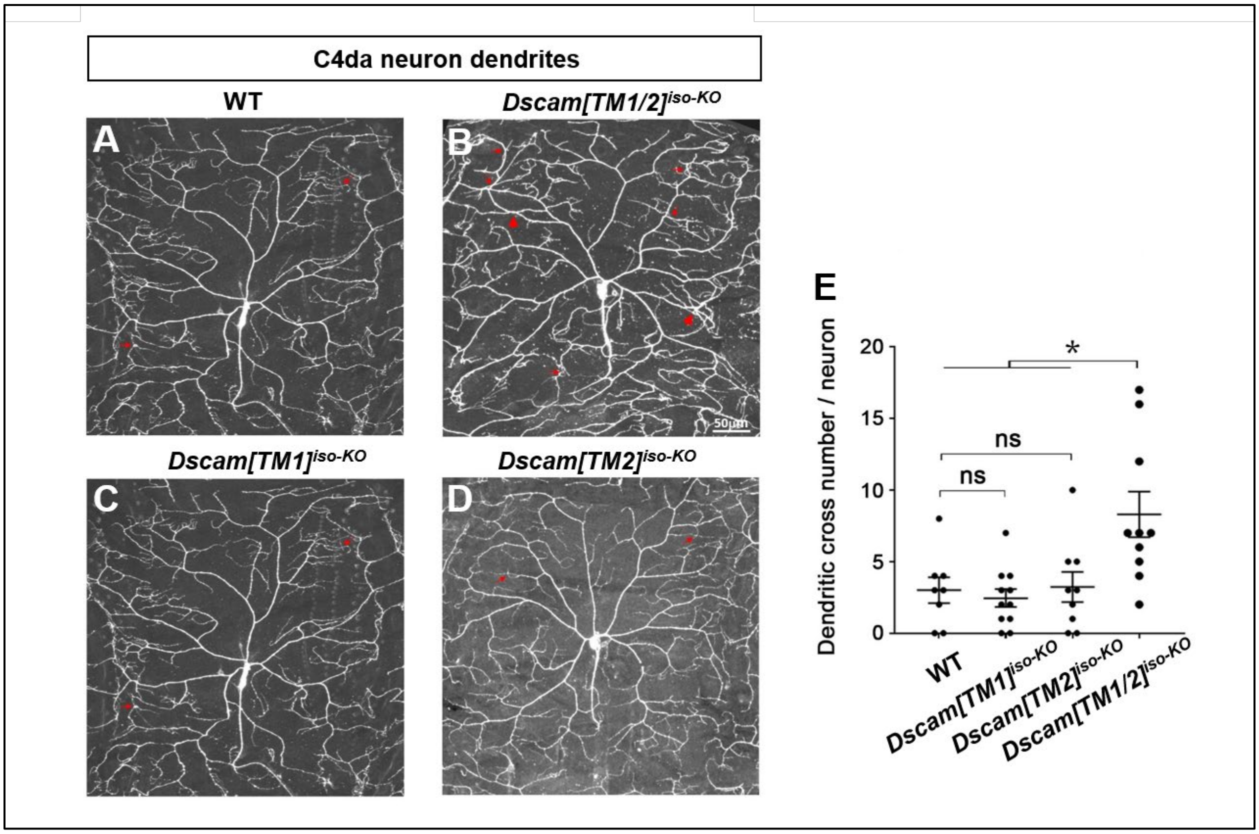


**Figure S4. *Dscam[TM1]* and *[TM2]* function redundantly in mediating dendritic self-avoidance in C4da neurons. (Related to Figure 3)**

(**A**-**D**) While *Dscam[TM1/2]*$^{iso\text{-}KO}$ (**B**) increases the crossing of dendritic branches in C4da neurons in 3$^{rd}$ instar larvae, neither *Dscam[TM1]*$^{iso\text{-}KO}$ nor *Dscam[TM2]*$^{iso\text{-}KO}$ impairs dendritic self-avoidance (**C** & **D**). Small red arrows point to crossings of fine dendritic branches, and large red arrows point to crossings of major dendritic branches, which is only observed when both isoforms are lost.

(**E**) Quantification of dendritic branch crosses in the C4da neuron ddaC.

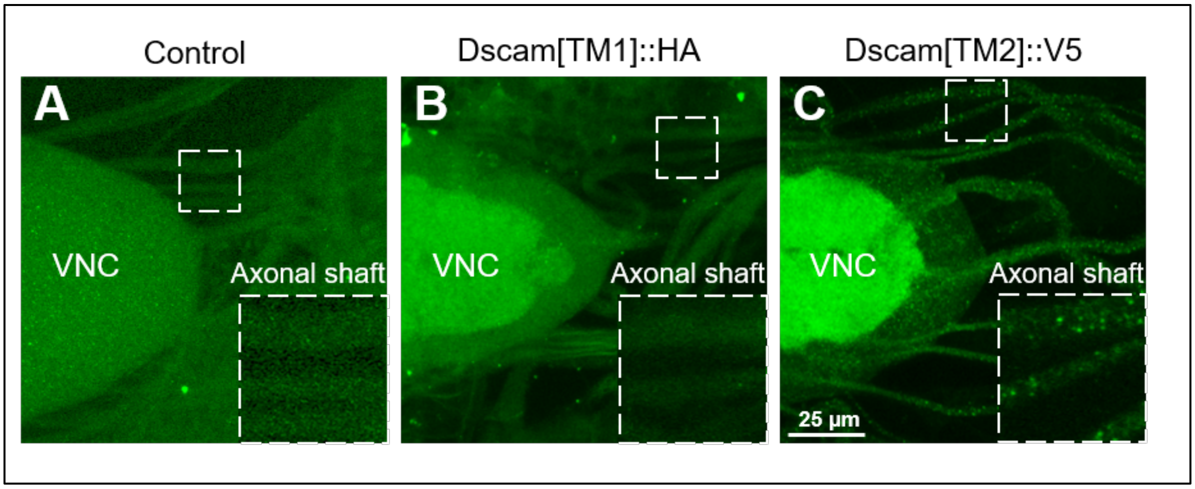


**Figure S5. Endogenous Dscam[TM2], but not [TM1], is localized in axons connecting the CNS and PNS. (related to Figure 4)**

Endogenous Dscam[TM1] is observed in VNC neuropil region, but not in axonal shafts. By contrast, endogenous Dscam[TM2] is localized in axon shafts besides the VNC neuropil. The insets are magnified views of the boxed regions.
